# Methane production by *Methanothrix thermoacetophila* via direct interspecies electron transfer with *Geobacter metallireducens*

**DOI:** 10.1101/2023.02.13.528421

**Authors:** Jinjie Zhou, Jessica A. Smith, Meng Li, Dawn E. Holmes

**Affiliations:** Archaeal Biology Center, Institute for Advanced Study, Shenzhen University, Shenzhen, Guangdong, China; Laboratory of Optoelectronic Devices and Systems of Ministry of Education and Guangdong Province, College of Optoelectronic Engineering, Shenzhen University, Shenzhen, Guangdong, China; Department of Microbiology, University of Massachusetts - Amherst, Amherst, Massachusetts, USA; Department of Biomolecular Sciences, Central Connecticut State University, New Britain, Connecticut, USA; Shenzhen Key Laboratory of Marine Microbiome Engineering, Institute for Advanced Study, Shenzhen University, Shenzhen, Guangdong, China; Department of Physical and Biological Science, Western New England University, Springfield, Massachusetts, USA

**Keywords:** *Methanothrix*, direct interspecies electron transfer (DIET), methane, magnetite, granular activated carbon (GAC), archaea, *Geobacter*, cytochromes, acetate

## Abstract

*Methanothrix* is widely distributed in natural and artificial anoxic environments and plays a major role in global methane emissions. It is one of only two genera that can form methane from acetate dismutation and through participation in direct interspecies electron transfer (DIET) with exoelectrogens. Although *Methanothrix* is a significant member of many methanogenic communities, little is known about its physiology. In this study, transcriptomics helped to identify potential routes of electron transfer during DIET between *Geobacter metallireducens* and *Methanothrix thermoacetophila*. Additions of magnetite to cultures significantly enhanced growth by acetoclastic methanogenesis and by DIET, while granular activated carbon (GAC) amendments impaired growth. Transcriptomics suggested that the OmaF-OmbF-OmcF porin complex and the octaheme outer membrane *c*-type cytochrome, Gmet_0930, were important for electron transport across the outer membrane of *G. metallireducens* during DIET with *Mx. thermoacetophila*. Clear differences in the metabolism of *Mx. thermoacetophila* when grown via DIET or acetate dismutation were not apparent. However, genes coding for proteins involved in carbon fixation and a surface associated quinoprotein, SqpA, were highly expressed in all conditions. Expression of gas vesicle genes was significantly lower in DIET-than acetate-grown cells, possibly to facilitate better contact between membrane associated redox proteins during DIET. These studies reveal potential electron transfer mechanisms utilized by both *Geobacter* and *Methanothrix* during DIET and provide important insights into the physiology of *Methanothrix* in anoxic environments.

**Importance:** *Methanothrix* is a significant methane producer in a variety of methanogenic environments including soils and sediments as well as anaerobic digesters. Its abundance in these anoxic environments has mostly been attributed to its high affinity for acetate and its ability to grow by acetoclastic methanogenesis. However, *Methanothrix* species can also generate methane by directly accepting electrons from exoelectrogenic bacteria through direct interspecies electron transfer (DIET). Methane production through DIET is likely to further increase their contribution to methane production in natural and artificial environments. Therefore, acquiring a better understanding of DIET with *Methanothrix* will help shedding light on ways to 1) minimize microbial methane production in natural terrestrial environments and 2) maximize biogas formation by anaerobic digesters treating waste.

## Introduction

*Methanothrix* (formerly *Methanosaeta*) species, arguably the most prodigious methanogens on earth, substantially contribute to the production of atmospheric methane and the conversion of wastes to methane biofuel (1). Species from this genus are frequently the most abundant methanogens in methanogenic terrestrial environments such as natural wetlands and flooded rice paddy soils and in many anaerobic digester systems (2-8). *Methanothrix* species are also important members of the methanogenic community in Arctic and Antarctic sediments (9-11), and scientists are particularly concerned with increases in methane production linked to melting permafrost in these polar terrestrial sediments as this will lead to a positive feedback loop that will further exacerbate climate change (12-14).

*Methanothrix* are one of only two methanogenic genera that can utilize acetate as a substrate for methanogenesis (1). Unlike the other acetoclastic methanogenic genus, *Methanosarcina, Methanothrix* have an extremely high affinity for acetate and generally maintain acetate at levels too low (μM range) for their competitors to metabolize (15). This ability to metabolize acetate at the low *in situ* levels found in most sediments and conventional mesophilic digesters is an important physiological capability, because acetate is a precursor for approximately two-thirds of the methane produced in terrestrial environments (1, 16) and is also an important intermediate in anaerobic digesters (17, 18).

Clearly the importance of acetate as a methane precursor demonstrates the central role of *Methanothrix* in carbon and electron flow in many methanogenic environments. However, it has also been found that *Methanothrix* species can reduce CO_2_ to methane by accepting electrons from exoelectrogenic bacteria via direct interspecies electron transfer (DIET) (4, 19), and this is likely to further increase their contribution to methane production in anoxic environments. Although *Methanothrix* species are predominant members of many methanogenic communities, little is known about their physiology (1, 20-23).

*Methanothrix* is considered a strictly acetoclastic methanogen, however, its genome contains genes from the CO_2_ reduction pathway which is used by hydrogenotrophic methanogens for growth on H_2_/CO_2_ and formate (1). Studies of *Methanothrix harundinacea* growing in co-culture with *Geobacter metallireducens* showed that it could convert ^14^C-labelled CO_2_ to ^14^C-methane, providing evidence that *Methanothrix* carries out CO_2_ reduction during DIET (4). In addition, CO_2_ reduction pathway genes were highly expressed by *Methanothrix* species growing by DIET in flooded rice paddy soils (2), anaerobic digesters treating waste (4, 24), and on the surface of cathodes in bioelectrochemical systems (BES) (25, 26). In a similar manner, acetoclastic *Methanosarcina* species growing by DIET with exoelectrogens highly expressed CO_2_ reduction genes (27-29).

In addition to *Methanothrix*’ high affinity for acetate (30), another feature that would make *Methanothrix* a strong competitor in low nutrient environments is the ability to fix CO_2_. All known *Methanothrix* species have genes coding for a RuBisCO-mediated reductive hexulose phosphate (RHP) pathway that forms formaldehyde as an intermediate (34, 35). They also have genes for a formaldehyde activating enzyme (Fae), an enzyme that catalyzes the conversion of formaldehyde into 5,10-Methylenetetrahydromethanopterin, an intermediate in the methanogenic CO_2_ reduction pathway. Metatranscriptomic data demonstrated that *Methanothrix* was highly expressing genes from the RHP pathway in an anaerobic digester (31).

In order to further understand the physiology of DIET between an exoelectrogen and *Methanothrix*, co-cultures were established between *G. metallireducens* and *Methanothrix thermoacetophila*. Previous studies have shown that conductive materials can enhance DIET between exoelectrogens and methanogens (32-37), and they are frequently added to bioreactors dominated by *Methanothrix* species (24). Therefore, to more fully understand the impact that conductive materials can have on *Methanothrix* physiology, two different conductive materials, granular activated carbon (GAC) and magnetite nanoparticles were added to *Mx. thermoacetophila* cultures. Transcriptomic studies were also done to identify potential pathways for electron transfer between the electron-donating (*G. metallireducens*) and electron-accepting (*Mx. thermoacetophila*) partners. These results provide valuable insights into *Methanothrix* physiology that can be used to better understand carbon flow in many methanogenic terrestrial ecosystems and to help optimize biomethane production from waste.

## Results and discussion

### *Methanothrix thermoacetophila* can accept electrons from *Geobacter metallireducens* via DIET

Co-cultures of *G. metallireducens* and *Mx. thermoacetophila* were grown with ethanol provided as the sole electron donor, and CO_2_ as the sole electron acceptor (Figure 1). Neither of the partners would survive alone in this medium, as the bacterium lacked an electron acceptor, and the methanogen lacked an electron donor. Utilization of H_2_/formate mediated electron transfer was also not possible because *G. metallireducens* does not produce H_2_ or formate during ethanol oxidation (38, 39), and *Mx. thermoacetophila* cannot use H_2_ or formate as an electron donor (40). Therefore, any methane produced by this co-culture resulted from DIET between *G. metallireducens* and *Mx. thermoacetophila*.

**Figure 1.**
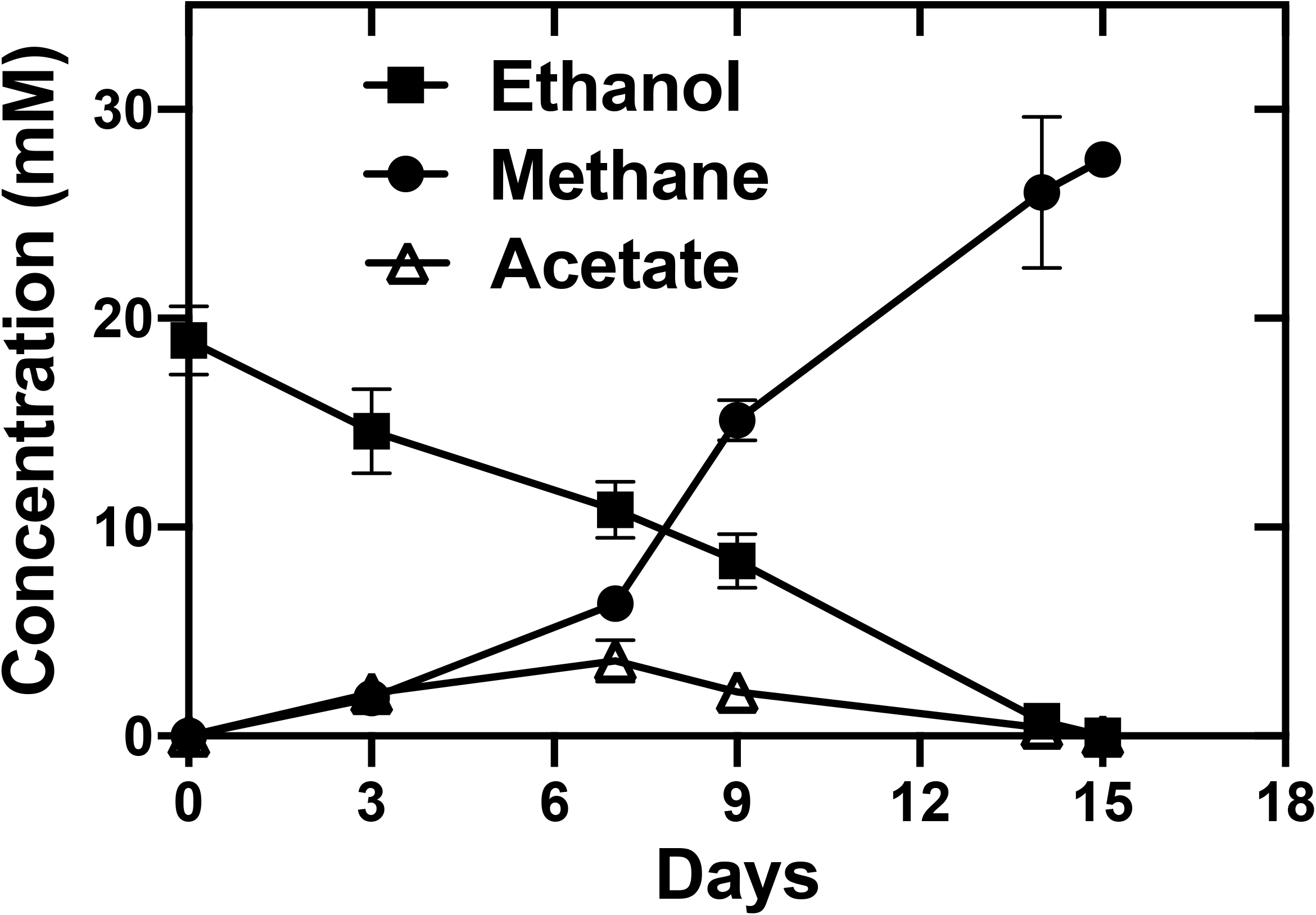
Growth of *G. metallireducens* and *Mx. thermoacetophila* co-cultures with ethanol (20 mM) provided as the sole electron donor and CO_2_ provided as the sole electron acceptor. Data represent means and standard deviations from triplicate cultures.

It took 95 days for initial co-culture aggregates to form, which is comparable to the time needed for co-cultures of *G. metallireducens* and *Mx. soehngenii* to become established (82 days) but much faster than co-cultures with *G. metallireducens* and *Mx. harundinacea* (167 days) (19) (Supplementary Figure S1). Once DIET was established, ethanol was converted to methane in 15 days with a yield of 1.46 mol methane/mol ethanol (Figure 1). This rate of methane production was significantly faster than the rate of established co-cultures of *G. metallireducens* and *Mx. harundinacea* which took 85 days to reduce 20 mM ethanol to methane (4). This rate was comparable to co-cultures established with *G. metallireducens* and *M. vacuolata* DH-1 (15 days), but was 1.9-2.3 times faster than rates seen in co-cultures with *G. metallireducens* and *Methanosarcina barkeri* (∼31 days), *Methanosarcina acetivorans* (28 days), or *Methanosarcina subterranea* (35 days) (28, 34, 41). Similar to other co-culture studies (4, 28, 34, 41), acetate concentrations initially increased until day 7 and then started to decline (Figure 1).

*G. metallireducens* and *Mx. thermoacetophila* formed visibly large, loosely clumped aggregates (Figure 2A), rather than the tight balls formed when *G. metallireducens* served as the electron-donating partner for DIET with other organisms (4, 39, 42). Confocal (Figure 2B, 2C) and TEM (Figure 2D-2G) images showed that *G. metallireducens* cells (short orange rods) were closely attached to the sheath of *Methanothrix* (green long filaments). Quantitative PCR of DNA extracted from these aggregates revealed that *G. metallireducens* accounted for over three quarters (76.03%± 2.18%) of the cells, likely because one long fiber-like cell of *Methanothrix* served as a scaffold for the attachment of multiple *Geobacter* cells.

**Figure 2.**
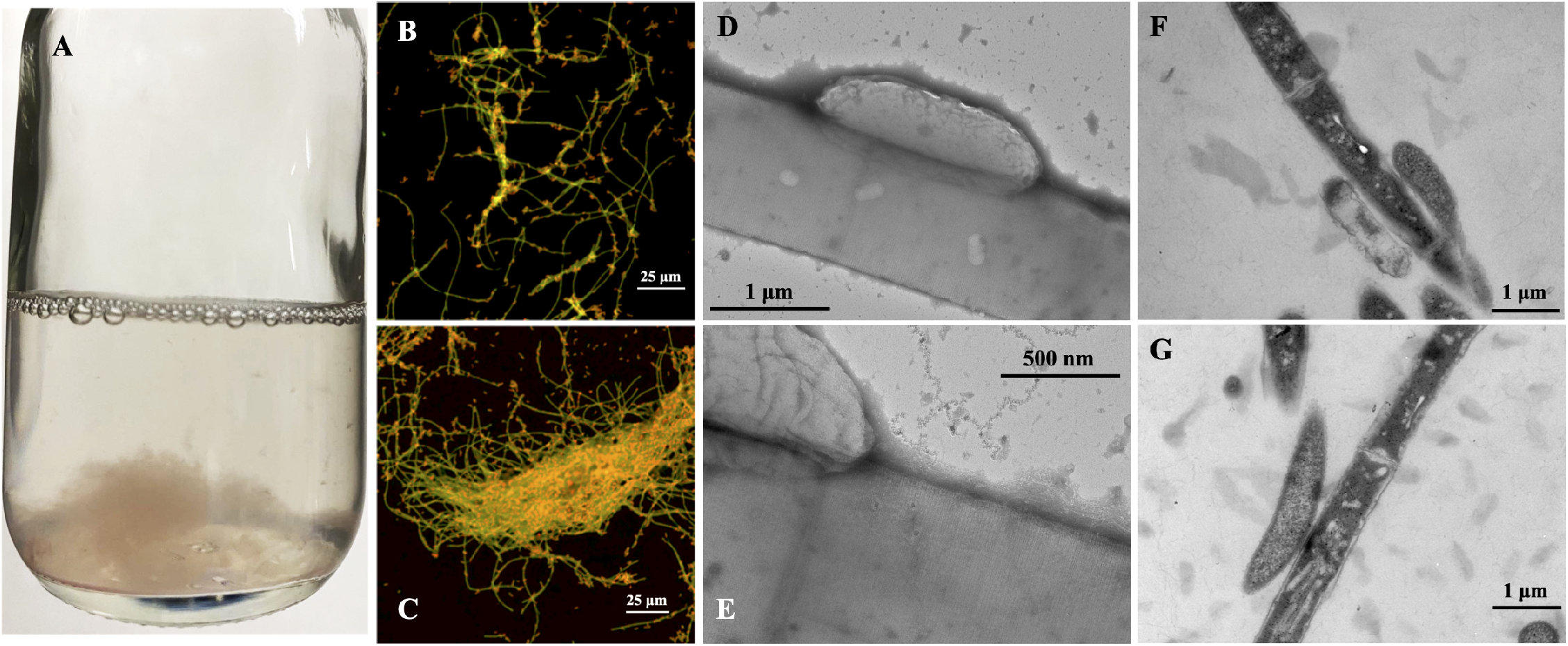
Morphology of *G. metallireducens* and *Mx. thermoacetophila* co-culture aggregates. (A) Appearance of loose aggregates visible to the naked eye; (B, C) FISH images showing the close attachment of *G. metallireducens* (short rod, orange) cells to *Mx. thermoacetophila* (long filament, green); (D, E) Negative-stain TEM images of co-cultures; (F, G) Ultrathin TEM images of co-cultures.

### Magnetite enhanced growth of *Mx. thermoacetophila*, while GAC had an inhibitory effect

Previous studies have shown that the addition of conductive materials such as granular activated carbon (GAC) (19, 37, 43) and magnetite (44) can stimulate DIET between *G. metallireducens* and an electron-accepting partner. Therefore, co-cultures were grown in the presence of both of these materials.

It was surprising to find that growth of *Mx. thermoacetophila* alone or in co-culture with *G. metallireducens* was severely impaired by the presence of GAC (Figure 3). In fact, acetoclastic methanogenesis was inhibited by GAC at concentrations above 20 g/L (Figure 3A). In addition, DIET co-cultures could not become established in the presence of GAC (Figure 3B, Supplementary Figure S1).

**Figure 3.**
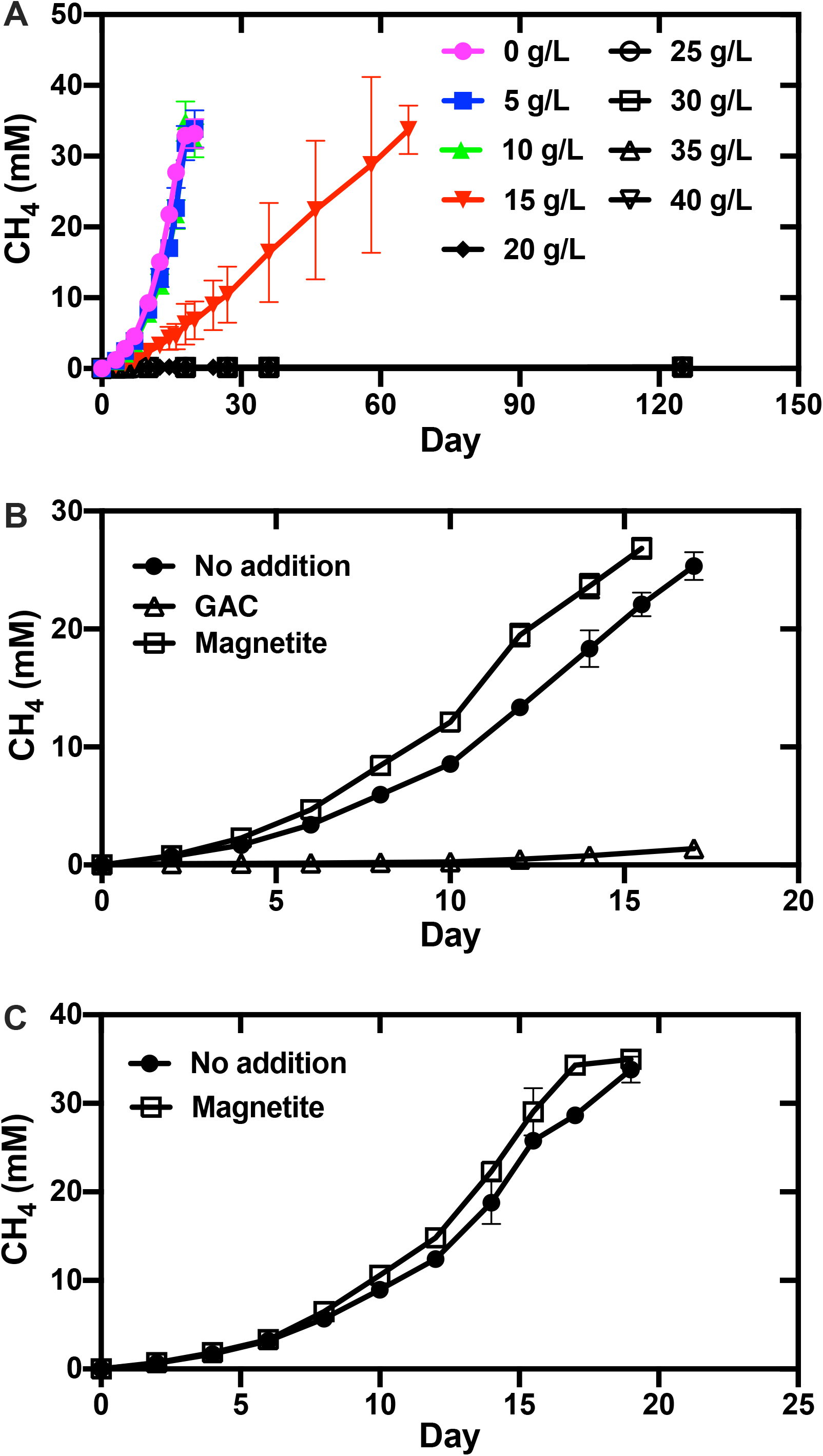
Effects of granular activated carbon (GAC) and magnetite on methane production by *Mx thermoacetophila*. (A) Pure cultures using acetate (40 mM) as substrate in the presence of various GAC concentrations; (B) co-cultures using ethanol (20 mM) as substrate in the presence of GAC (40 g/L) or magnetite (10 mM); (C) pure cultures using acetate (40 mM) as substrate in the presence of magnetite (10 mM). Data represent the average and standard deviations from triplicate cultures.

GAC is frequently added to anaerobic digesters to promote methane production (45, 46), and in many cases, the proportion of *Methanothrix* species declines while *Methanosarcina* species are enriched in GAC-amended reactors (47-50). The results from this study help explain why *Methanothrix* is often less abundant in GAC-amended reactors and indicate that GAC may not be the best option for stimulation of methanogenesis in environments where *Methanothrix* is an important member of the community.

Similar to previous studies with *Methanosarcina* (51, 52), the addition of magnetite slightly stimulated (1.2 times faster; *p*-value = 0.05) the rate of growth for pure cultures grown by acetoclastic methanogenesis (Figure 3C). Growth in co-culture was also enhanced and aggregates only took 45 days to form compared to 95 days in non-amended co-cultures (Supplementary Figure S1). Once aggregates became established, they grew at rates that were 1.2 times faster (*p*-value = 0.007) (Figure 3B). Consistent with these results, magnetite additions have also been shown to stimulate methanogenesis by *Methanothrix* species participating in DIET in anaerobic digesters (53) and by *Methanosarcina* growing with *Geobacter* in methanogenic rice paddy soil enrichments (54).

### The *G. metallireducens* DIET transcriptome

#### *G. metallireducens* transcriptomes are similar when cells are grown in co-culture with *Mx. thermoacetophila* or *Methanosarcina barkeri*

The *G. metallireducens* transcriptome during growth by DIET with *Mx. thermoacetophila* (MX) was compared to its DIET transcriptome when grown with other electron-accepting partners (*M. barkeri* (MB), *M. acetivorans* (MA), *M. subterranea* (MS), and *Geobacter sulfurreducens* (GS)) (55). Multidimensional scaling analysis plots (MDS plots) generated with the biological coefficient of variation (BCV) method (Figure 4) revealed that the transcriptome of *G. metallireducens* grown in co-culture with *Mx. thermoacetophila* was most similar to its transcriptome when it was grown in co-culture with the Type I *Methanosarcina* species, *M. barkeri* (41). Similar to *M. barkeri, Mx. thermoacetophila* lacks outer surface *c*-type cytochromes and does not have any potentially conductive archaella (28, 41). Transcriptomes from *G. metallireducens* cells grown by DIET with electron-accepting partners with cytochromes and potentially conductive e-pili/archaella (*G. sulfurreducens* and the Type II *Methanosarcina* species *M. acetivorans* and *M. subterranea*) were significantly different.

**Figure 4.**
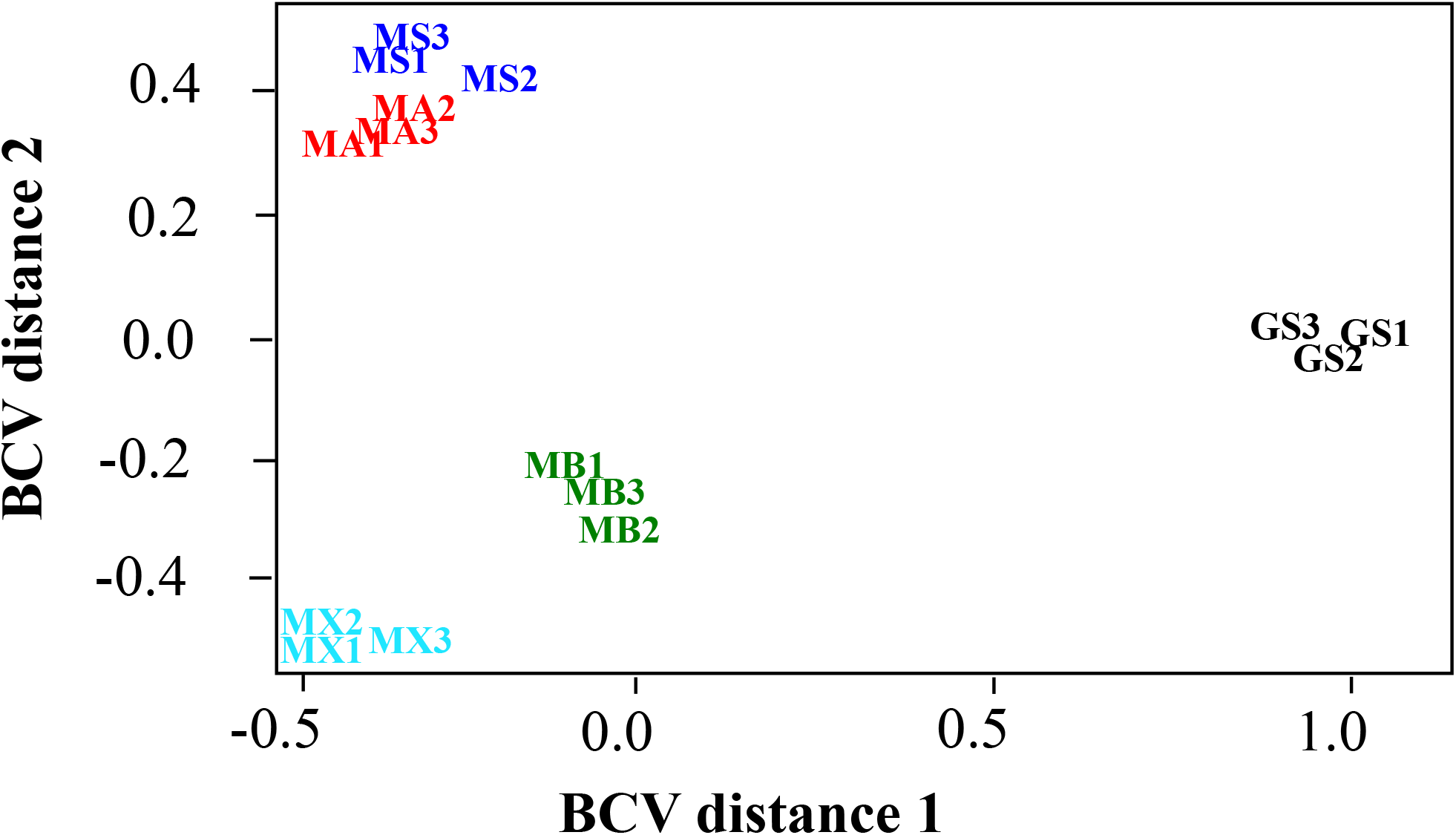
Comparison of *G. metallireducens* RNAseq libraries from co-cultures with *Mx. thermoacetophila* (MX), *Methanosarcina barkeri* (MB), *Methanosarcina acetivorans* (MA), *Methanosarcina subterranea* (MS), and *Geobacter sulfurreducens* (GS) using multidimensional scaling analysis with the biological coefficient of variation (BCV) method.

### Genes from the PccF porin-cytochrome complex were more highly expressed in DIET-grown cells

Electron transport across the outer membrane of *Geobacter* cells to extracellular electron acceptors such as insoluble Fe(III) oxides or other microorganisms requires porin-*c*-type-cytochrome (Pcc) complexes (56, 57). *G. metallireducens* has three Pcc complexes; PccF (Gmet_0908-0910), PccG (Gmet_0911-0913), and PccH (Gmet_0825-0827) (55, 58). Similar to previous studies of DIET with *G. metallireducens* serving as the electron-donating partner (55), genes from the PccH complex were not highly expressed by *G. metallireducens* cells grown in co-culture with *Mx. thermoacetophila*. However, the number of transcripts from PccF complex genes, *omcF* (Gmet_0910), *omaF* (Gmet_0909), and *ombF* (Gmet_0908) were 6.0 (*p*-value = 1.4×10^−11^), 5.3 (*p*-value = 7.2×10^−12^), and 5.9 (*p*-value = 4.1×10^−11^) times more abundant in DIET-grown cells than Fe(III) respiring cells (Table 1 and Supplementary Table S1). Levels of transcripts from PccG complex genes, *omcG* (Gmet_0913), *omaG* (Gmet_0912), and *ombG* (Gmet_0911) were lower than the median RPKM values (Supplementary Table S2) indicating that PccG was not important for DIET with *Mx. thermoacetophila*. Addition of magnetite to the *G. metallireducens*/*Mx. thermoacetophila* co-cultures increased expression of PccG gene transcripts (Table 1), but the expression was still well below the median RPKM value (Supplementary Table S2). This same pattern of increased expression of PccF relative to PccG genes was observed when *G. metallireducens* was grown in co-culture with *M. barkeri* (55).

**Table 1.** Differences in expression of genes coding for electron transfer proteins in *G. metallireducens* cells that were grown under various conditions. Values represent fold differences between *G. metallireducens* cells grown by DIET with *Mx. thermoacetophila* compared to growth with ethanol (20 mM) as the electron donor and Fe(III) citrate (56 mM) as the electron acceptor (DIET *vs*. Fe(III)-citrate); *G. metallireducens* cells grown by DIET with *Mx. thermoacetophila* in the presence of 10 mM magnetite *vs*. DIET without magnetite (DIET-magnetite *vs*. DIET); or *G. metallireducens* cells grown by DIET with *Mx. thermoacetophila* without magnetite compared to DIET with magnetite (DIET *vs*. DIET-magnetite). Only genes with fold differences >2 and *p*-values <0.05 were considered significant. NS: no significant difference. *p*-values are available in Supplementary Table S1.

| Locus ID | Annotation | Gene | Location | DIET vs.<br>Fe(III)-<br>citrate | DIET-magnetite vs.<br>DIET | DIET vs.<br>DIET-magnetite |
| --- | --- | --- | --- | --- | --- | --- |
| Gmet_0910 | 10 heme *c*-type cytochrome from PccF complex | *omcF* | Outer membrane/surface | 6.00 | 2.96 | NS |
| Gmet_0909 | 9 heme periplasmic *c*-type cytochrome from PccF complex | *omaF* | Periplasmic | 5.32 | 2.76 | NS |
| Gmet_0908 | Porin protein from the PccF complex | *ombF* | Outer membrane/surface | 5.88 | 2.11 | NS |
| Gmet_0913 | 9 heme outer membrane *c*-type cytochrome from PccG | *omcG* | Outer membrane/surface | NS | 2.78 | NS |
| Gmet_0912 | 8 heme periplasmic *c*-type cytochrome from PccG complex | *omaG* | Periplasmic | NS | 4.79 | NS |
| Gmet_0911 | Porin protein from the PccG complex | *ombG* | Outer membrane/surface | NS | 3.64 | NS |
| Gmet_1399 | Type IV major pilin subunit, PilA | *pilA* | Outer membrane/surface | NS | NS | 2.96 |
| Gmet_1400 | Short pilin chaperone protein, Spc | *spc* | Outer membrane/surface | NS | NS | 4.29 |
| Gmet_2928 | 7 heme *c*-type cytochrome protein from CbcABCDE | *cbcA* | Periplasmic | 28.86 | NS | NS |
| Gmet_2929 | *b*-type cytochrome from CbcABCDE complex | *cbcB* | Inner membrane | 41.70 | NS | NS |
| Gmet_2930 | Cytochrome *c* family protein, 11 hemes | *cbcC* | Periplasmic | 2.55 | NS | NS |
| Gmet_2931 | Mono-heme *c*-type cytochrome protein from CbcABCDE complex | *cbcD* | Periplasmic | 2.00 | NS | NS |
| Gmet_2932 | Membrane protein from CbcABCDE complex | *cbcE* | Inner membrane | 2.80 | 2.35 | NS |
| Gmet_0533 | Membrane protein from CbcMNOPQR | *cbcQ* | Inner membrane | 51.56 | NS | NS |
| Gmet_0534 | *c*-type cytochrome protein, 5 hemes | *cbcR* | Periplasmic | 28.19 | NS | NS |
| Gmet_0930 | *c*-type cytochrome, 8 hemes |  | Outer membrane/surface | 20.45 | NS | NS |
| Gmet_1210 | *c*-type cytochrome, 2 hemes | *ccpB* | Periplasmic | 5.95 | NS | NS |
| **Gmet_0252** | MonoHEME *c*-type cytochrome protein | *coxB* | Inner membrane | 4.53 | NS | NS |
| **Gmet_1809** | *c*-type cytochrome, 5 hemes | *actA* | Inner membrane | 3.98 | NS | NS |
| **Gmet_1019** | *c*-type cytochrome, 2 hemes | *narC* | Inner membrane | 3.81 | NS | NS |
| **Gmet_2626** | MonoHEME P460 *c*-type cytochrome |  | Unknown | 3.37 | NS | NS |
| **Gmet_0294** | *c*-type cytochrome, 4 hemes | *nrfA* | Periplasmic | 3.14 | NS | NS |
| **Gmet_3091** | *c*-type cytochrome protein, 2 hemes | *macA* | Periplasmic | 3.07 | 2.69 | NS |
| **Gmet_0679** | *c*-type cytochrome, 5 hemes |  | Unknown | 2.84 | NS | NS |
| **Gmet_1197** | *c*-type cytochrome, 5 hemes |  | Unknown | 2.71 | 3.14 | NS |
| **Gmet_2930** | 11 heme *c*-type cytochrome protein, CbcABCD complex | *cbcC* | Periplasmic | 2.55 | NS | NS |
| **Gmet_0142** | *c*-type cytochrome, 8 hemes |  | Periplasmic | 2.44 | NS | NS |
| **Gmet_1814** | MonoHEME *c*-type cytochrome | *actE* | Unknown | 2.30 | NS | NS |
| **Gmet_0292** | Diheme *c*-type cytochrome |  | Outer membrane/surface | 2.08 | NS | NS |
| **Gmet_1647** | Diheme *c*-type cytochrome |  | Unknown | 2.07 | NS | NS |
| **Gmet_0558** | *c*-type cytochrome protein, 27 hemes | *omcO* | Unknown | NS | 10.62 | NS |
| **Gmet_1087** | MonoHEME *c*-type cytochrome |  | Unknown | NS | 7.19 | NS |
| **Gmet_0571** | *c*-type cytochrome protein, 34 hemes |  | Outer membrane/surface | NS | 3.23 | NS |
| **Gmet_0121** | Diheme *c*-type cytochrome |  | Unknown | NS | 2.89 | NS |
| **Gmet_2896** | *c*-type cytochrome protein, 4 hemes |  | Outer membrane/surface | NS | 2.79 | NS |
| **Gmet_0600** | *c*-type cytochrome protein, 19 hemes |  | Outer membrane/surface | NS | 2.67 | NS |
| **Gmet_2839** | *c*-type cytochrome protein, 11 hemes |  | Periplasmic | NS | 2.65 | NS |
| **Gmet_1094** | Diheme *c*-type cytochrome | *cccA* | Unknown | NS | 2.45 | NS |
| **Gmet_0330** | MonoHEME *c*-type cytochrome | *narH* | Inner membrane | NS | 2.36 | NS |
| **Gmet_0557** | *c*-type cytochrome protein, 4 hemes |  | Unknown | NS | 2.25 | NS |
| **Gmet_0598** | *c*-type cytochrome protein, 22 hemes |  | Outer membrane/surface | NS | 2.15 | NS |
| **Gmet_1846** | Diheme *c*-type cytochrome | *ppcE* | Periplasmic | NS | NS | 3.06 |
| **Gmet_2902** | Diheme *c*-type cytochrome | *ppcA* | Periplasmic | NS | NS | 4.08 |
| **Gmet_3166** | Diheme *c*-type cytochrome | *ppcB* | Periplasmic | NS | NS | 2.75 |
| **Gmet_1867** | *c*-type cytochrome protein, 8 hemes |  | Unknown | NS | NS | 2.45 |
| **Gmet_2432** | Monoheme P460 *c*-type cytochrome |  | Unknown | NS | NS | 5.57 |

### Pilin genes were not more highly expressed in DIET-grown cells

Although genes coding for PilA, the conductive monomer of e-pili (59), and Spc, the putative e-pili chaperone protein (60), were being expressed by DIET-grown cells at levels >3.5 times above the median RPKM values (Supplementary Table S2), significant differences in expression were not observed between DIET- and Fe(III)-respiring cells (Table 1). These results suggest that e-pili may not be as important for DIET with *Mx. thermoacetophila* as they are for DIET with *M. acetivorans* and *G. sulfurreducens*, both of which required e-pili for DIET-based growth and expressed *pilA* and *spc* genes at levels that were >2 fold higher in DIET grown cells than Fe(III) respiring cells (28, 38). Other electron-accepting species lacking *c*-type cytochromes, *M. barkeri* and *Methanobacterium electrotrophus* do not require e-pili for DIET (36, 55, 61). Therefore, *Mx. thermoacetophila* may be another electron-accepting partner lacking outer surface *c*-type cytochromes that does not require e-pili for participation in DIET. However, further studies with gene deletion strains are required.

It was also interesting to find that the addition of magnetite to the co-cultures decreased the expression of e-pili associated genes; *pilA* (Gmet_1399) and *spc* (Gmet_1400) transcripts were 3.0 (*p*-value = 2.3×10^−9^) and 4.3 (*p*-value = 8.6×10^−11^) times less abundant in DIET co-cultures amended with magnetite (Table 1, Supplementary Table S1). Studies have indicated that magnetite can compensate for pilin associated *c*-type cytochromes involved in extracellular electron exchange during DIET between *Geobacter* species (44, 62). However, evidence that magnetite can substitute for electrically conductive pili during electron transfer between *Geobacter* and *Methanothrix* is currently not available.

### Genes coding for Gmet_0930 and the CbcABCDE complex were highly expressed by DIET-grown cells

Genes coding for two periplasmic multiheme *c*-type cytochromes (Gmet_2928 and Gmet_0534) and an outer surface multiheme *c*-type cytochrome (Gmet_0930) were >20 times more highly expressed by *G. metallireducens* cells grown by DIET with *Mx. thermoacetophila* than Fe(III)-respiring cells (Table 1). Gmet_2928 codes for CbcA, a periplasmic 7-heme cytochrome that is part of the cytochrome *bc* complex CbcABCDE (Gmet_2928-2932). Cytochrome *bc* (Cbc) complexes are composed of a transmembrane *b*-type cytochrome in close association with at least one multiheme *c*-type cytochrome, and they are involved in shuttling electrons from the quinone pool in the inner membrane to *c*-type cytochromes found in the periplasmic space (57, 63, 64). The gene coding for CbcA and other components of this complex, the *b*-type cytochrome (CbcB), two other periplasmic *c*-type cytochromes (CbcC and CbcD), and a membrane protein (CbcE), were 28.9 (*p*-value = 2.0×10^−14^), 41.7 (*p*-value = 5.7×10^−14^), 2.6 (*p*-value = 2.2 × 10^−6^), 2.0 (*p*-value = 2.1×10^−6^), and 2.8 (*p*-value = 1.5×10^−8^) times more highly expressed in DIET-grown cells than Fe(III)-respiring cells (Table 1, Supplementary Table S1).

Gmet_0534 codes for a 5-heme periplasmic cytochrome putatively associated with another Cbc complex, CbcMNOPQR (Gmet_0533-0539). Although this gene was 28.2 times (*p*-value = 3.5×10^−14^) more highly expressed in DIET-grown cells, *cbcO* was the only other gene from this complex that was more highly expressed by DIET-grown cells (Table 1, Supplementary Table S1). Genes from both the CbcABCDE and CbcMNOPQR complexes were highly expressed by *G. metallireducens* cells grown by DIET with other electron-accepting partners and during growth with insoluble Fe(III) oxide, however, genetic studies showed that they were not required for growth in either of these conditions (55, 65).

Gmet_0930 encodes an 8-heme outer surface *c*-type cytochrome that was 20.4 (*p*-value = 7.5×10^−14^) times more highly expressed by DIET-grown than Fe(III)-respiring cells (Table 1, Supplementary Table S1). *G. metallireducens* requires this *c*-type cytochrome for formation of DIET aggregates with *G. sulfurreducens* (55) and it was among the most highly expressed genes by *G. metallireducens* cells grown in co-culture with *M. acetivorans, M. subterranea*, and *M. barkeri*. Although Gmet_0930 deletion mutant strains eventually adapted to grow in co-culture with all three *Methanosarcina* species, growth was impaired even after 4 transfers indicating that Gmet_0930 was also important for DIET with these other partners (55).

Significant differences in expression of the periplasmic cytochrome PpcA (Gmet_2902) were not observed between DIET grown and Fe(III)-respiring cells. However, PpcA (Gmet_2902) was the second most highly expressed *c*-type cytochrome with values that were 21 times higher (*p*-values = 3.2×10^−5^) than median RPKM values for both DIET and DIET-magnetite conditions (Supplementary Table S2). PpcA is highly conserved among *Geobacter* species and it is required for optimal Fe(III) reduction by *G. sulfurreducens* (66, 67). Based on analysis of transcriptomic data, it is proposed that the route for electron transfer from the quinone pool in the *G. metallireducens* inner membrane to *Mx. thermoacetophila* during DIET requires activity from the CbcABCDE complex, periplasmic PpcA, the PccF conduit (*omaF, ombF*, and *omcF*), and the outer surface *c*-type cytochrome Gmet_0930 (Figure 5). Electrons may then be transferred directly to *Mx. thermoacetophila* or they may first be transferred to e-pili.

**Figure 5.**
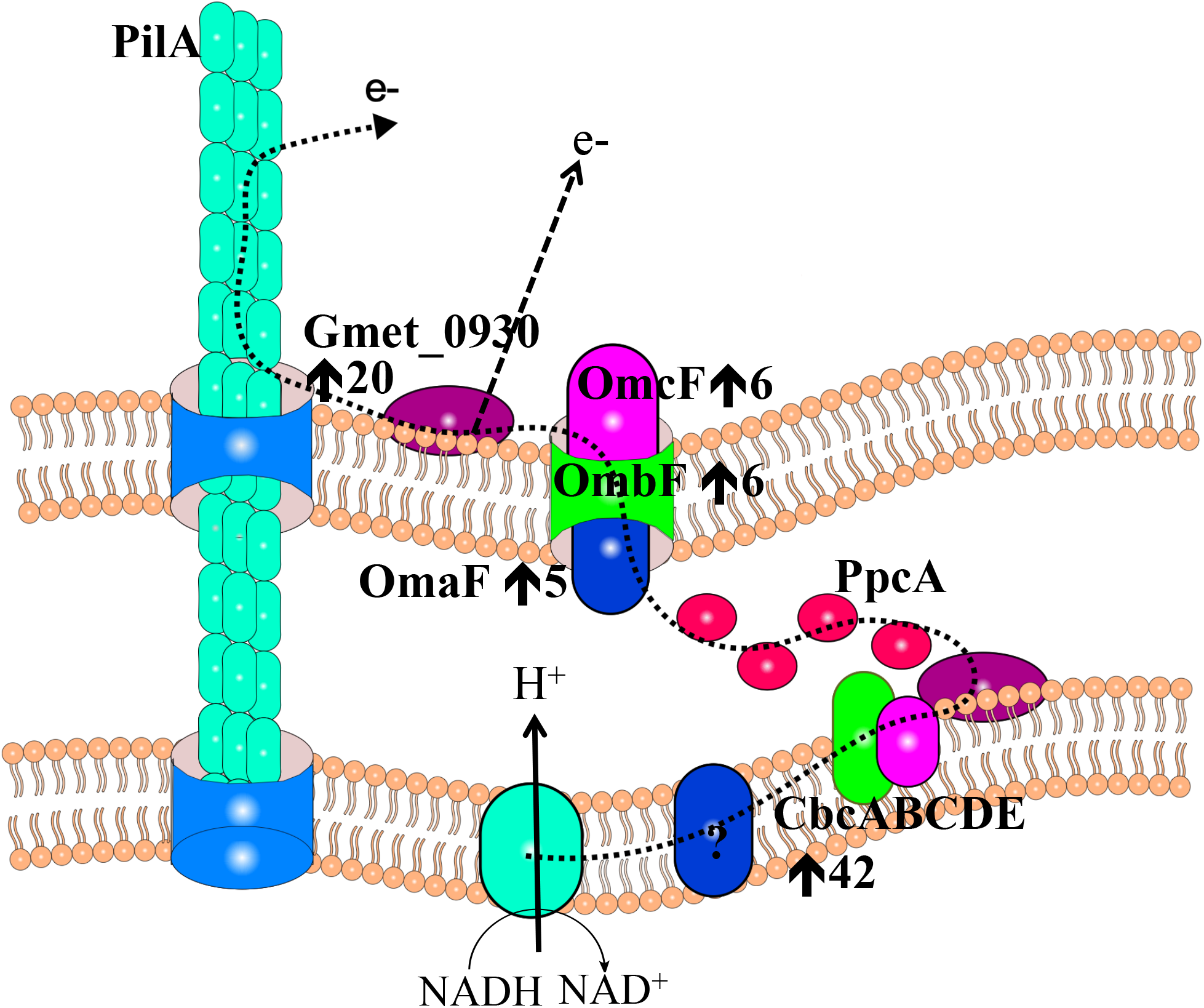
Pathway proposed for electron transfer from *G. metallireducens* to *Mx. thermoacetophila* during DIET. Electrons are transferred from the quinone pool in the inner membrane to the CbcABCDE quinone-oxidoreductase complex, then to the periplasmic *c*-type cytochrome PpcA, which then shuttles electrons to the PccF (OmaF, OmbF, and OmcF) porin-cytochrome complex and then to the outer surface octaheme cytochrome Gmet_0930. Electrons may then be transferred directly to *Mx. thermoacetophila* or they may first be transferred to conductive e-pili, which can then interact with the *Mx. thermoacetophila* surface. Arrows represent fold up-regulated in DIET grown cells compared to cells grown with ethanol as the electron donor and Fe(III) citrate as the electron acceptor (For proteins composed of multiple subunits, values from the most highly expressed subunits are shown). If an arrow is not listed with a protein from the proposed pathway, the gene was not differentially expressed by DIET-or Fe(III) respiring cells.

### The *Mx. thermoacetophila* DIET transcriptome

Unfortunately, analysis of transcriptomic data did not reveal many significant differences between DIET-based and acetoclastic metabolism in *Mx. thermoacetophila*. In all of the conditions, most genes from pathways for methanogenesis from acetate and CO_2_, carbon fixation, the reductive citric acid cycle, and carbon monoxide and formate metabolism had RPKM values that were more than 2 fold above median RPKM values (Supplementary text, Supplementary Figure S2, Supplementary Table S3). Many of these genes were also more highly expressed under both acetoclastic and DIET-based conditions in the presence of magnetite.

More specifically, comparison of DIET- to acetate-grown cells revealed that genes from the CO_2_ reduction and the RHP carbon fixation pathways were being highly expressed by cells in both conditions. Previous studies have suggested that these genes are only highly expressed by DIET-grown *Methanothrix* cells (2, 4, 25, 31). However, transcriptomic comparisons between DIET-grown and acetoclastic cells were not done in these experiments.

The only clear differences in expression patterns between DIET- and acetate-grown cells were related to genes coding for gas vesicle proteins. Two gene clusters (Mthe_0055-0063 and Mthe_0069-0073) coding for gas vesicle proteins were >2 times (*p*-value < 0.05) more highly expressed in acetate-grown cells than DIET-grown cells in the presence and absence of magnetite (Table 2). Gas vesicles are commonly found in archaeal sheaths, including those from *Mx. thermoacetophila*, and make cells more buoyant within a water column (68, 69). This decrease in expression of gas vesicles during DIET may facilitate better contact between redox proteins on the surface of *G. metallireducens* and *Mx. thermoacetophila*.

**Table 2.** Differences in expression of genes coding for gas vesicle proteins in *Mx. thermoacetophila* cells grown under various conditions. Values represent fold differences between *Mx. thermoacetophila* cells grown by acetoclastic methanogenesis *vs*. cells grown by DIET with *G. metallireducens* (Acetate *vs*. DIET) as well as in the presence of 10 mM magnetite (Acetate-magnetite *vs*. DIET magnetite); *Mx. thermoacetophila* cells grown by DIET with *G. metallireducens* without magnetite compared to DIET with magnetite (DIET *vs*. DIET-magnetite); *Mx. thermoacetophila* cells grown by acetoclastic methanogenesis compared to acetoclastic methanogenesis with magnetite (Acetate *vs*. Acetate-magnetite). NS: no significant difference. All *p*-values <0.05 and are available in Supplementary Table S4.

| Locus ID | Annotation | Gene | Acetate vs. DIET | Acetate-magnetite vs. DIET-magnetite | DIET vs. DIET-magnetite | Acetate vs. Acetate-magnetite |
| --- | --- | --- | --- | --- | --- | --- |
| **Mthe_0055** | Gas vesicle synthesis GvpLGvpF fusion | *gvpLF* | 2.19 | 1.72 | 2.62 | 3.34 |
| **Mthe_0056** | Gas vesicle protein GvpO | *gvpO* | 2.19 | 1.79 | 1.96 | 2.46 |
| **Mthe_0058** | Gas vesicle protein GvpN | *gvpN* | 2.27 | 2.05 | 1.71 | 1.88 |
| **Mthe_0060** | Gas vesicle synthesis protein GvpA | *gvpA* | 2.68 | 2.17 | 3.96 | 5.48 |
| **Mthe_0061** | Gas vesicle synthesis protein GvpA | *gvpA* | 2.63 | 2.17 | 4.12 | 5.02 |
| **Mthe_0062** | Gas vesicle synthesis protein GvpA | *gvpA* | 2.63 | 2.17 | 4.12 | 5.02 |
| **Mthe_0063** | Gas vesicle synthesis protein GvpA | *gvpA* | 2.63 | 2.17 | 4.12 | 5.02 |
| **Mthe_0069** | Gas vesicle protein GvpG | *gvpG* | 2.94 | NS | 4.53 | 11.07 |
| **Mthe_0070** | Gas vesicle protein GvpA | *gvpA* | 2.89 | NS | 1.94 | 6.25 |
| **Mthe_0071** | Gas vesicle protein GvpA | *gvpM* | 2.63 | NS | 2.82 | 8.85 |
| **Mthe_0072** | Gas vesicle protein GvpK | *gvpK* | 2.69 | NS | 2.40 | 5.71 |
| **Mthe_0073** | Gas vesicle synthesis GvpLGvpF fusion | *gvpLF* | 2.47 | NS | 2.06 | 4.72 |

Expression of *gvp* genes was also lower in DIET- and acetate-cells grown in the presence of magnetite. When the medium was supplemented with 10 mM magnetite, *Mx. thermoacetophila* sheaths were coated with magnetite particles (Supplementary Figure S3). It is possible that cells do not need to produce as many gas vesicles in the presence of magnetite for several reasons; 1) magnetite can act as an electron conductor during DIET, 2) the magnetite particles can serve as a scaffold for biofilm formation during DIET or acetoclastic growth, and 3) acetate can adsorb to the positively charged magnetite particles reducing the need for cells to maintain buoyancy in the medium. Although transcriptomic data suggest that reduced expression of Gvp proteins is advantageous for DIET or growth in the presence of magnetite, differences in gas vesicle abundance were not obvious in negative-stain and ultrathin TEM images (data not shown).

### Possible routes for electron uptake by *Mx. thermoacetophila*

As discussed above, there were clear differences in expression of many genes in the presence or absence of magnetite (Supplementary Table S4). However, these differences were not apparent in genes coding for surface associated proteins that could potentially facilitate direct electron uptake from *G. metallireducens* (Table 3). This suggests that these genes may not be differentially regulated when cells are grown under varying conditions. Analysis of the most highly expressed surface proteins led us to the following proposed routes for electron uptake into *Mx. thermoacetophila*. However, this analysis is still speculative.

**Table 3.**
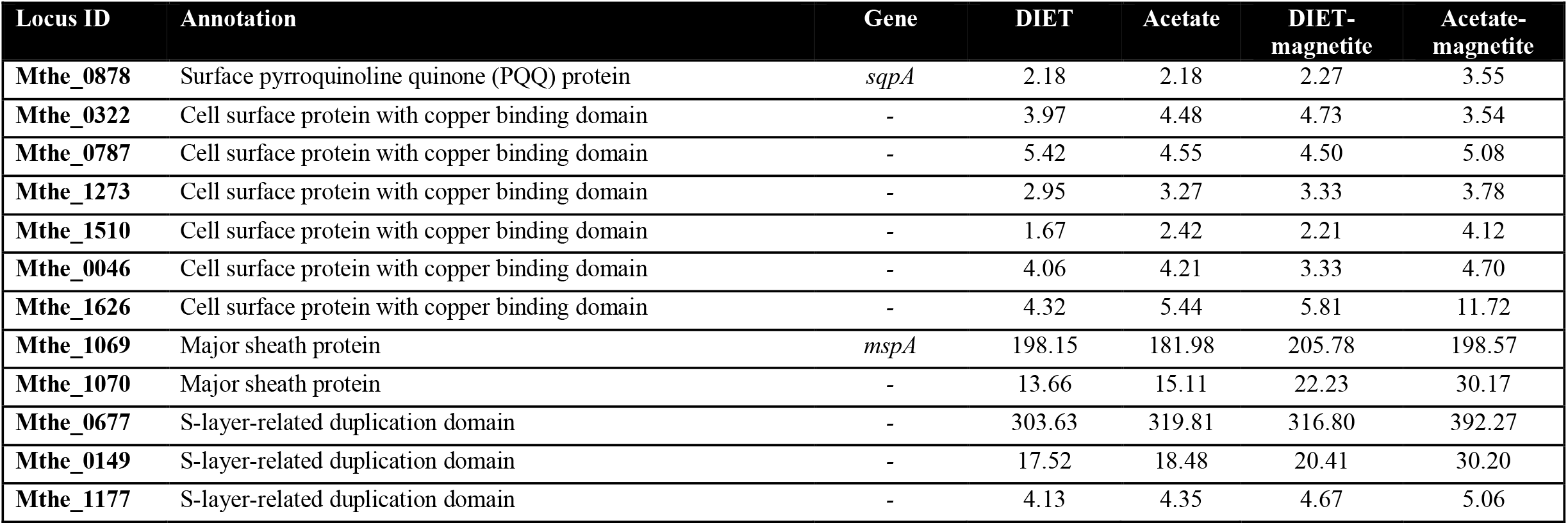
Fold differences from median RPKM values for *Mx. thermoacetophila* genes coding for surface proteins that could potentially facilitate electron uptake from *G. metallireducens* in all 4 conditions. **DIET**: *Mx. thermoacetophila* cells grown in co-culture with *G. metallireducens* with ethanol (20 mM) as the electron donor (median log_2_ RPKM value was 7.31), **Acetate**: *Mx. thermoacetophila* grown with acetate (40 mM) (median log_2_ RPKM value was 7.30), **DIET-magnetite**: DIET with *Mx. thermoacetophila* and *G. metallireducens* with ethanol (20 mM) as the electron donor in the presence of magnetite (10 mM) (median log_2_ RPKM value was 7.29), **Acetate-magnetite**: *Mx. thermoacetophila* grown with acetate (40 mM) in the presence of magnetite (10 mM) (median log_2_ RPKM value was 7.27). All *p*-values are <0.05 and were calculated by ANOVA using the R statistical package. *p*-values are available in Supplementary Table S3.

The main sheath fiber protein (MspA; Mthe_1069) was the most highly expressed surface-associated gene in all of the conditions and had >181 times (*p*-values < 1.14×10^−5^) higher expression than median RPKM values (Table 3, Supplementary Figure S2; Supplementary Table S3). The MspA sheath protein forms amyloid fibrils with extended beta-sheet structures (68) and numerous amino acid residues (10.7% of the protein residues; Supplementary Figure S4). Previous studies have shown that stacked aromatic amino acids in amyloid fibrils confer conductivity (70), suggesting that MspA could be conductive.

Unlike *Geobacter* or Type II *Methanosarcina* species (41, 57, 71, 72), *Mx. thermoacetophila* does not have any surface multiheme *c*-type cytochromes that could readily accept electrons from an extracellular electron donor (1, 73). However, the genome does have a gene coding for a transmembrane pyrroloquinoline quinone (PQQ) binding protein; surface quinoprotein A (*sqpA*; Mthe_0878) that was highly expressed (Table 3). The PQQ binding site of quinoproteins has a beta-propeller fold composed of antiparallel β-sheets radially arranged around a central tunnel (74, 75). The mature SqpA protein has an extremely high concentration of aromatic amino acids (11.3%) (Supplementary Figure S4), and pi stacking of aromatic residues arranged in the center of the funnel-shaped propeller could enhance electron transfer properties of this redox protein. Evidence that surface quinoproteins could be involved in DIET comes from observations that *M. barkeri* was expressing genes encoding surface quinoproteins during growth via DIET with *G. metallireducens* (27) and *Rhodopseudomonas palustris* (29).

It is possible that electrons accepted by a membrane associated electron carrier such as SqpA are being funneled to Fpo dehydrogenase to reduce F_420_ or ferredoxin in a manner similar to that suggested by *Methanosarcina* (Supplementary Figure S2) (27-29). Unlike *mspA* and *sqpA*, Fpo dehydrogenase genes were differentially expressed in cells grown in the presence of magnetite (Supplementary Table S3). In addition, elucidation of the role that Fpo dehydrogenase might play in DIET is further complicated by the fact that it is also likely to be involved in acetoclastic methanogenesis (21). The Fpo dehydrogenase complex of *Mx. thermoacetophila* lacks FpoF, which is the subunit that accepts electrons from F_420_H_2_ in *Methanosarcina* species (23, 76). Rather, it has been proposed that iron clusters in the FpoB or FpoI subunits can accept electrons from reduced ferredoxin during acetoclastic growth, and heterodisulfide reduction was observed when *Mx. thermoacetophila* membranes were incubated with reduced ferredoxin from *Methanosarcina mazei* (21).

Another possibility is that a soluble iron-sulfur flavoprotein from the same family as FpoF (77, 78) interacts with the Fpo dehydrogenase complex to reduce F_420_. The *Mx. thermoacetophila* genome has two genes (Mthe_0174 and Mthe_0959) from the FpoF family that code for proteins that are most similar to the beta subunit from the F_420_ hydrogenase complex, FrhB. These proteins are not likely to be part of a hydrogenase complex, as the genome lacks genes that code for the other two subunits from this complex (FrhA and FrhG) (79), and hydrogenase activity has not been detected in *Mx. thermoacetophila* cells (21). One of these FrhB genes (Mthe_0174) was expressed at levels that were > 6.6 times (*p*-values < 2.6×10^−6^) higher than the median RPKM values in all conditions (Supplementary Table S3).

Reduced ferredoxin and/or F_420_H_2_ generated by Fpo dehydrogenase could then transfer electrons to either the soluble heterodisulfide reductase complex, HdrABC, or the membrane-bound heterodisulfide complex, HdrDE, to reduce CoM-S-S-CoB (Supplementary Figure S2). Levels of transcripts for *hdrA* (Mthe_1576) and *hdrDE* (Mthe_0980-0981) were > 7.4 times (*p*-values < 1.1×10^−6^) and > 15.4 times (*p*-values < 2.7×10^−5^) higher than the median RPKM values for all conditions (Supplementary Table S3).

## Conclusions

*Geobacter metallireducens* and *Methanothrix thermoacetophila* were able to grow syntrophically by coupling the oxidation of ethanol with the reduction of CO_2_ to methane. Addition of the conductive material, magnetite enhanced methanogenesis by acetate dismutation and by DIET, while GAC amendments impaired growth. Transcriptomic studies revealed that *G. metallireducens* uses mechanisms for electron transport to *Mx. thermoacetophila* that are similar to those used for electron transport to the Type I *Methanosarcina, M. barkeri*. Both *M. barkeri* and *Mx. thermoacetophila* lack outer surface multiheme *c*-type cytochromes and putatively conductive archaella. Transcription of genes coding for gas vesicle proteins was down-regulated during DIET or in the presence of magnetite likely because buoyancy within the water column is not required when cells can adhere to a surface. These results provide invaluable insight into *Methanothrix* physiology. However, further studies are required to fully understand the role of *Methanothrix* in methanogenic ecosystems.

## Materials and Methods

### Culture conditions

*Methanothrix thermoacetophila* DSM 6194 was cultured anaerobically (N_2_/CO_2_ 80/20, v/v) at 55°C in DSMZ 334 medium (https://www.dsmz.de/microorganisms/medium/pdf/DSMZ_Medium334.pdf) with acetate (40 mM) as the substrate. *Geobacter metallireducens* GS-15 (ATCC 53774) was routinely cultured anaerobically (N_2_/CO_2_ 80/20, v/v) at 30°C in freshwater medium with ethanol (20 mM) as the electron donor and ferric citrate (56 mM) as the electron acceptor (80).

It was necessary to adapt both *G. metallireducens* and *Mx. thermoacetophila* to similar growth conditions before co-culture experiments could be conducted. *G. metallireducens* was adapted to grow in *Methanothrix* medium (DSMZ 334), in which acetate and sulfide were omitted, and ethanol (20 mM) and ferric citrate (56 mM) were supplied as the electron donor and the electron acceptor, respectively. The cultivation temperature for *G. metallireducens* was gradually increased from 30 to 42°C, while the cultivation temperature for *Mx. thermoacetophila* was gradually decreased from 55 to 42°C.

After *G. metallireducens* and *Mx. thermoacetophila* were adapted to grow at 42°C, an equal proportion (10%) of the two partners was added to modified DMSZ 334 medium with ethanol (20 mM) as the electron donor and CO_2_ as the electron acceptor. Sulfide (0.5 mM) and L-cysteine·HCl (1 mM) were added from sterile anoxic stocks. The co-cultures were cultivated anaerobically (N_2_/CO_2_ 80/20, v/v) at 42°C.

When noted, granular activated carbon (GAC, Sigma, C2889, various concentrations) or magnetite nanoparticle (10 mM) was added to the medium before autoclaving. Magnetite was prepared as previously described (44).

### RNA extraction and transcriptome analyses

Triplicate cultures (co-cultures, pure cultures of *Mx. thermoacetophila* and pure cultures of *G. metallireducens*) were harvested during the mid-logarithmic phase for transcriptomic analyses. Specifically, cells from co-cultures and pure cultures of *Mx. thermoacetophila* were collected when methane concentrations reached ∼18 mM, and *G. metallireducens* cells were collected when Fe(II) concentrations were ∼35 mM. Pellets from all samples were formed by centrifugation in 50 mL conical tubes at 4000 × *g* for 15 min at 4°C. After centrifugation, the pellets were frozen in liquid nitrogen and stored at - 80°C until RNA extraction procedures were performed.

Total RNA from sample pellets was extracted as previously described (81). Whole mRNAseq libraries were generated using the NEB Next® UltraTM Directional RNA Library Prep Kit for Illumina® (New England Biolabs, MA, USA). The clustering of index-coded samples was then performed on a cBot Cluster Generation System. After cluster generation, the library was sequenced on an Illumina Novaseq6000 platform and 150 bp paired-end reads were generated (Magigene Biotechnology, Guangzhou, China).

Raw data was checked with FASTQC (http://www.bioinformatics.babraham.ac.uk/projects/fastqc/), trimmed with Trimmomatic (82), and merged with FLASH (83). Ribosomal RNA (rRNA) reads were then removed from the libraries with SortMeRNA (84). The trimmed mRNA reads were mapped against genomes of *G. metallireducens* (CP000148) and *Mx. thermoacetophila* (CP000477) using SeqMan NGen (DNAStar). Reads were then normalized and processed for differential expression studies using the edgeR package in Bioconductor (85). All genes that were ≥1.5-fold differentially expressed with *p*-values of ≤0.05 are reported in Supplementary Tables.

### DNA extraction and quantitative PCR

Genomic DNA was extracted from triplicate co-cultures with the MasterPure complete DNA purification kit (Lucigen). The proportion of *G. metallireducens* and *Mx. thermoacetophila* cells in co-cultures was determined with quantitative PCR using the following primer pairs: (i) Gm-f (5’-ATGGCCCACATCTTCATCTC -3’) and Gm-r (5’-TGCATGTTTTCATCCACGAT-3’) which amplified a 104 bp fragment from the *bamY* gene (Gmet_2143) encoding benzoate-CoA ligase of *G. metallireducens* (86), and (ii) Mx-f (5’-GAGGATCTTGCCCGGATATT-3’) and Mx-r (5’-TATTGTAACGCCAGAGCCTC-3’) which amplified a 102 bp fragment from the *sseA* gene (Mthe_1071) encoding the rhodanese domain protein of *Mx. thermoacetophila*. Quantitative PCR was performed with iTaq Universal SYBR Green Supermix (Bio-Rad) on a QuantStudio 3 Real-Time PCR system (Applied Biosystems).

### Microscopy

Microbial cells were routinely checked with phase-contrast and fluorescence microscopy (Nikon E600) to ensure that cultures were not contaminated. Fluorescence *in situ* hybridization (FISH) of cell aggregates was conducted as previously described with a few modifications (87). Briefly, co-culture cells were fixed with 2% paraformaldehyde and 0.5% glutaraldehyde in 50 mM PIPES (pH 7.2) at 4°C for 2 h, followed by dehydration in 70% ethanol for 30 min. Cells were then transferred to glass slides, air dried, and immersed in hybridization buffer (900 mM NaCl, 20 mM Tris, 10% formamide, 0.01% SDS, 5 ng/μL each of the probes, pH 7.2) at 46°C for 2h. Next, the slides were washed in washing buffer (450 mM NaCl, 20 mM Tris-HCl, 5 mM EDTA, 0.01% SDS, pH 7.2) at 48°C for 30 min, rinsed gently with Milli-Q water and examined with a laser scanning confocal microscope (Nikon Eclipse Ti2). Probes used in this study were MX825 (5’-[cy5]-TCGCACCGTGGCCGACACCTAGC) for *Mx. thermoacetophila* and Geo1 (5’-[cy3]-AGAATCCAAGGACTCCGT) for *G. metallireducens* (4).

For negative-stained transmission electron microscopy (TEM), cells of *Mx. thermoacetophila* from either pure cultures or co-cultures were deposited on carbon-coated copper grids (200-mesh) for 10 min and stained with 2% phosphotungstic acid for 1 min. The grids were then air-dried and examined with a transmission electron microscope (ITACHI-HT7700) at 80 kV. For ultrathin TEM, the aggregates of cocultures and pure cultures of *Mx. thermoacetophila* were fixed with 2% paraformaldehyde and 2.5% glutaraldehyde in 100 mM phosphate buffer (pH 7.2) at 4°C overnight and embedded in low-melt agarose (1.5% in phosphate buffer). The agarose-embedded aggregates were then fixed with 1% osmium tetroxide for 3 h, dehydrated in gradient ethanol solution (30%, 50%, 70%, 90%, 95%, and 100% twice), embedded in Spi-pon 812 resin, polymerized, sectioned, stained with lead citrate and examined with a transmission electron microscope (HITACHI-HT7800) at 80 kV.

For scanning electron microscopy (SEM), cells of *Mx. thermoacetophila* collected during the mid-logarithmic phase were fixed with 2.5% glutaraldehyde in 100 mM phosphate buffer (pH 7.2) at 4°C overnight. Cells were then washed with phosphate buffer three times and post-fixed with 1% osmium tetroxide for 1 h. Fixed cells were dehydrated at 4°C with gradient ethanol solution (30%, 50%, 70%, 80%, 90%, 95%, and 100% twice) for 20 min for each step. Cells were then treated with ethanol/isoamyl acetate (v/v=1:1) for 30 min, and pure isoamyl acetate overnight. After dehydration, the samples were dried with a critical point dryer, coated with gold, and observed under a scanning electron microscope (HITACHI-SU8010) at 3 kV.

### Analytical techniques

Ethanol concentrations were measured with a gas chromatograph equipped with a flame ionization detector (Clarus 600; PerkinElmer Inc., CA). Acetate concentrations were measured by high-performance liquid chromatography (SHIMADZU, Japan) with an Aminex HPX-87H Ion Exclusion column (300 mm × 7.8 mm) and an eluent of 8.0 mM sulfuric acid. Methane was monitored by gas chromatography with a flame ionization detector (SHIMADZU, GC-8A) (88). Ferrous iron concentrations were determined by first incubating cultures in 0.5 N HCl and then measuring Fe(II) concentrations with a ferrozine assay at an absorbance of 562 nm as previously described (89).

## Data availability

Illumina sequence reads have been submitted to the Sequence Read Archive (SRA) of the NCBI database under BioProject PRJNA914893 and Biosamples SAMN32360990-32360994.

## Acknowledgments

We thank the Instrument Analysis Center of Shenzhen University for the assistance with the collection of transmission electron microscopy images.

This research was supported by the National Natural Science Foundation of China (42207144, 32225003, 92251306, and 31970105), the China Postdoctoral Science Foundation (2021TQ0212), the Shenzhen Science and Technology Program (JCYJ20200109105010363), and the Innovation Team Project of Universities in Guangdong Province (2020KCXTD023).

## Ethics declarations

The authors do not declare any conflicts of interest.

